# A near-complete dataset of plant growth form, life history, and woodiness for all Australian plants

**DOI:** 10.1101/2023.12.19.572473

**Authors:** Elizabeth Wenk, David Coleman, Rachael Gallagher, Daniel Falster

**Affiliations:** Evolution and Ecology Research Centre, University of New South Wales Sydney, Sydney, Australia; Hawkesbury Institute for the Environment, Western Sydney University, Sydney, Australia

## Abstract

Tabular records of plant trait data are essential for diverse research purposes. Here we present scorings for a trio of core plant traits, plant growth form, woodiness, and life history, for nearly all accepted taxon concepts included in the Australian Plant Census (APC). This dataset is predominately derived from Australia’s state and national floras, supplemented by the taxonomic literature and diverse web resources. In total, 29,993 species and infraspecific taxa were scored for plant growth form, 30,279 for woodiness, and 30,056 for life history, with taxa scored as displaying a single or multiple trait values, as appropriate. This resource will enable rapid assessment of plant responses to disturbance events and new biogeographic analyses of trait distributions.

## Introduction

To provide large-scale descriptions of plant biodiversity, information on trait values must exist alongside taxon lists, occurrence data, and phylogenies. Combined, this infrastructure informs us about what species exist, where they are found (occurrences), and their form and function. Incomplete trait-by-taxon matrices hinder our ability to rapidly assess functional diversity or include traits in descriptions of biodiversity or as covariates in analyses. Data on species occurrences are now readily available through GBIF (at a global scale) or the Atlas of Living Australia (ALA) for Australia-specific analyses, but the limited knowledge of the functional traits associated with each taxon introduces biases into large-scale biogeographic analyses (Andrew et al. 2021). Meanwhile, managers are unable to rapidly predict or assess species’ responses to major disturbance events if their functional traits are not readily available in a tabular format. For instance, growth form is a core trait often documented to distinguish between broad types of plants, and plants with these different core life histories may exhibit contrasting responses to climatic shifts or disturbance events (Butt and Gallagher 2018; Gallagher et al. 2021). Ultimately, despite a strong interest in plant traits and morphological and functional strategies, the patchy coverage of tabular trait data across national (or global) floras has limited the scope and uptake of trait-based approaches, especially at large spatial scales.

Recent years have seen ongoing efforts to expand coverage of a few core traits. Although standardised global compilations are beginning to emerge (Weigelt, König, and Kreft 2020; Díaz et al. 2022; Taylor et al. 2023), data on Australian plants are underrepresented in global trait compilations (Maitner et al. 2023), furthering the importance of developing and maintaining compilations dedicated to the Australian flora . AusTraits is a large compilation of trait data specifically for Australian plants, including data on plant growth form for 23,338 taxa, on life history for 22,031 taxa, and on woodiness of 9,026 taxa in its first public release (v3.0.2, 14 July 2021; Falster et al. 2021). When released in 2021, it offered a substantial improvement on other databases for Australia, but coverage still fell far short of providing trait values for all 30,490 species and infraspecific taxa in the Australian flora.

Although data do not yet exist for most traits across most taxa, taxonomic descriptions in their raw form—or aggregated into floras—contain a wealth of trait data and span most taxa. Basic morphological trait data is available for most described species and some of this content links directly to ecological function, such as plant growth form and woodiness (Coleman et al. 2023). In addition, nearly all taxon descriptions include the taxon’s life history. For the Australian flora, taxon descriptions have been compiled into six mostly complete state and territory floras and a partially complete national flora. There also exist regional compilations, such as the Flora of Southeast Queensland. Combined with the original taxonomic treatments, these contain near-complete coverage of these fundamental ecological traits, just not in a structured tabular format.

Recently, Coleman et al. (2023) used human-guided text processing to extract trait values from the text paragraphs given as taxon descriptions in floras. This effort greatly increased coverage for plant growth form, woodiness and life history, as well as a diversity of morphological traits such as leaf dimensions, leaf shape, fruit type, and plant storage organs. These data are available through the AusTraits trait database, v4.2, 18 September 2023. However, the outputs of the automated workflow remained incomplete and required manual reconciling.

In this paper, we aim to further extend Coleman et al. (2023)’s compilation by:

1. Gap-filling values from original taxonomic descriptions,
2. Manually reviewing ambiguous trait values in the textual paragraphs in state and national floras, referencing additional resources as required, and
3. Reconciling alternative values from different sources into a harmonised value for each taxon.

Together, these provide a near-complete single resource for 30,279 taxa or 99.3% of accepted taxa in the Australian Plant Census, or plant growth form, life history, and woodiness.

## Methods and Results

We focussed on extending and refining tabular data compilations for three traits—plant growth form, life history and woodiness—for all taxon concepts accepted by the Australian Plant Census (APC). The APC (https://biodiversity.org.au/nsl/services/search/taxonomy) offers a list of the accepted taxon concepts for all Australian vascular plants, including both native and naturalised taxa. As of November 2022, the APC included 30,494 accepted taxa of rank species, subspecies, varieties, and forms. Trait definitions were sourced from the AusTraits Plant Dictionary (APD; Table 1). The APD is a recent effort to formalise and publish definitions for all traits concepts used by AusTraits and includes allowed values for each categorical trait (APD; Wenk et al. 2023). The curated values include both broad, near-universal descriptions of plant growth form, life history and woodiness, as well as some more nuanced terms describing the unique forms and functions of Australia’s plants. These definitions were the basis for trait values extracted both here and by Coleman et al. (2023).

**Table 1.**
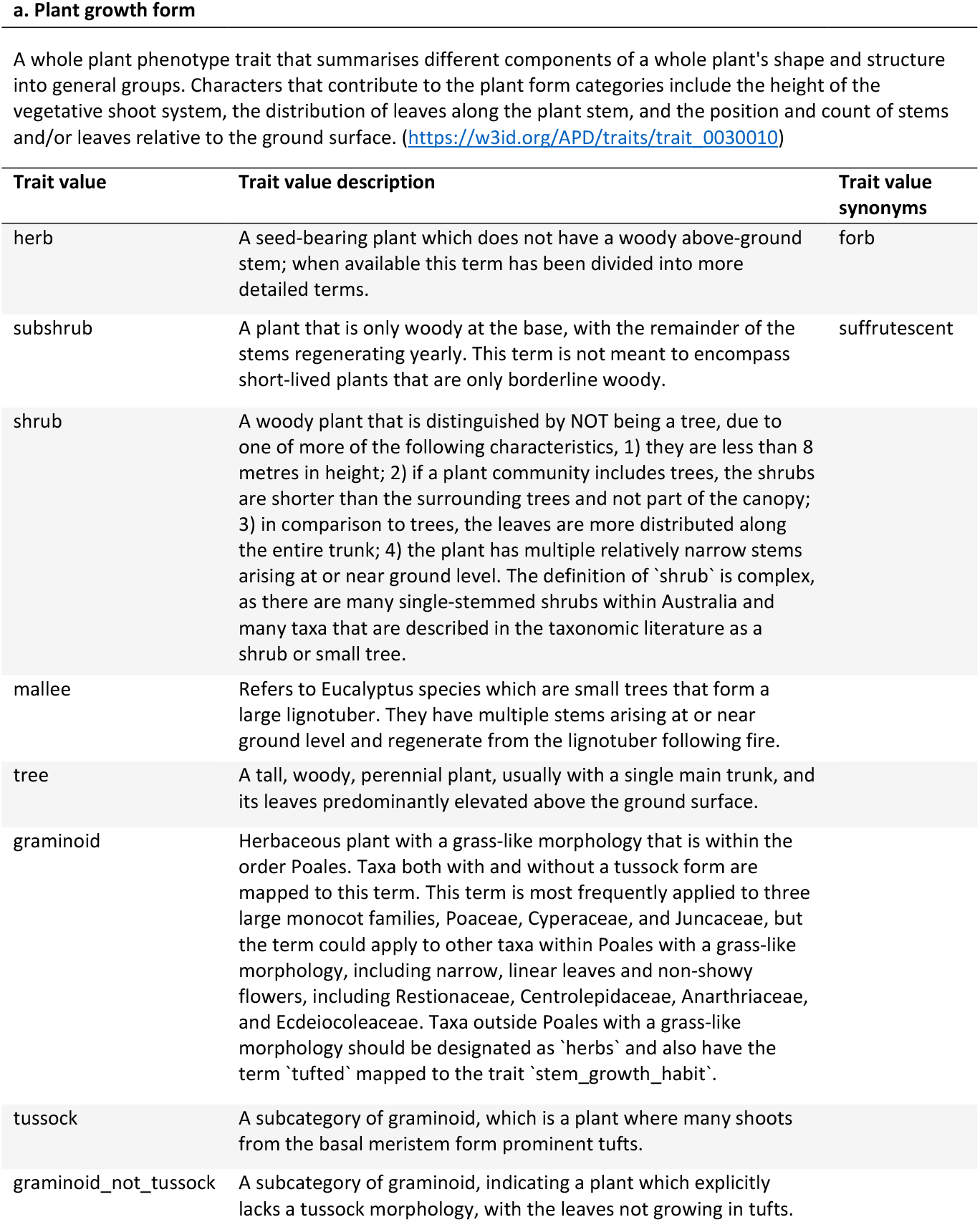
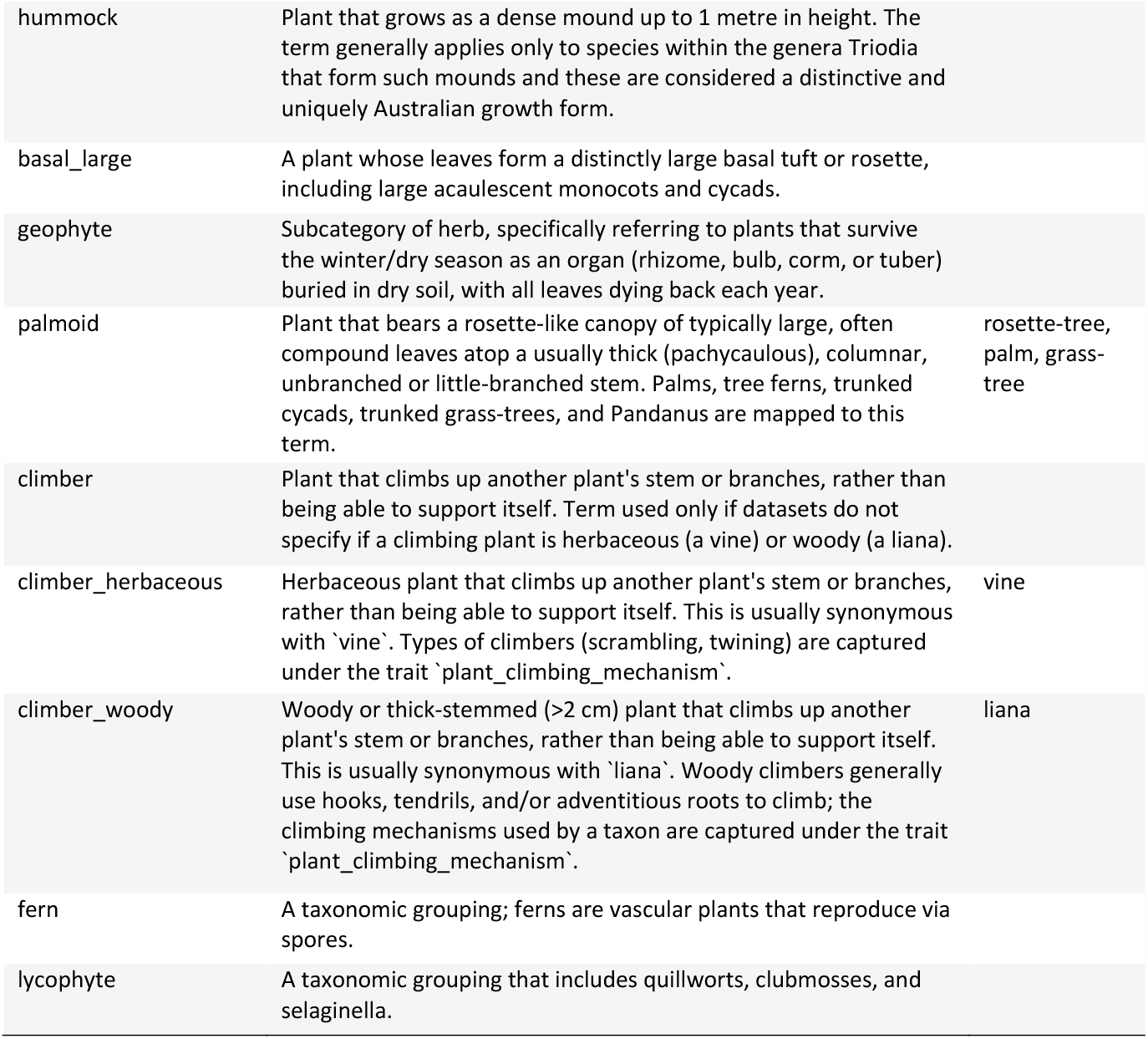
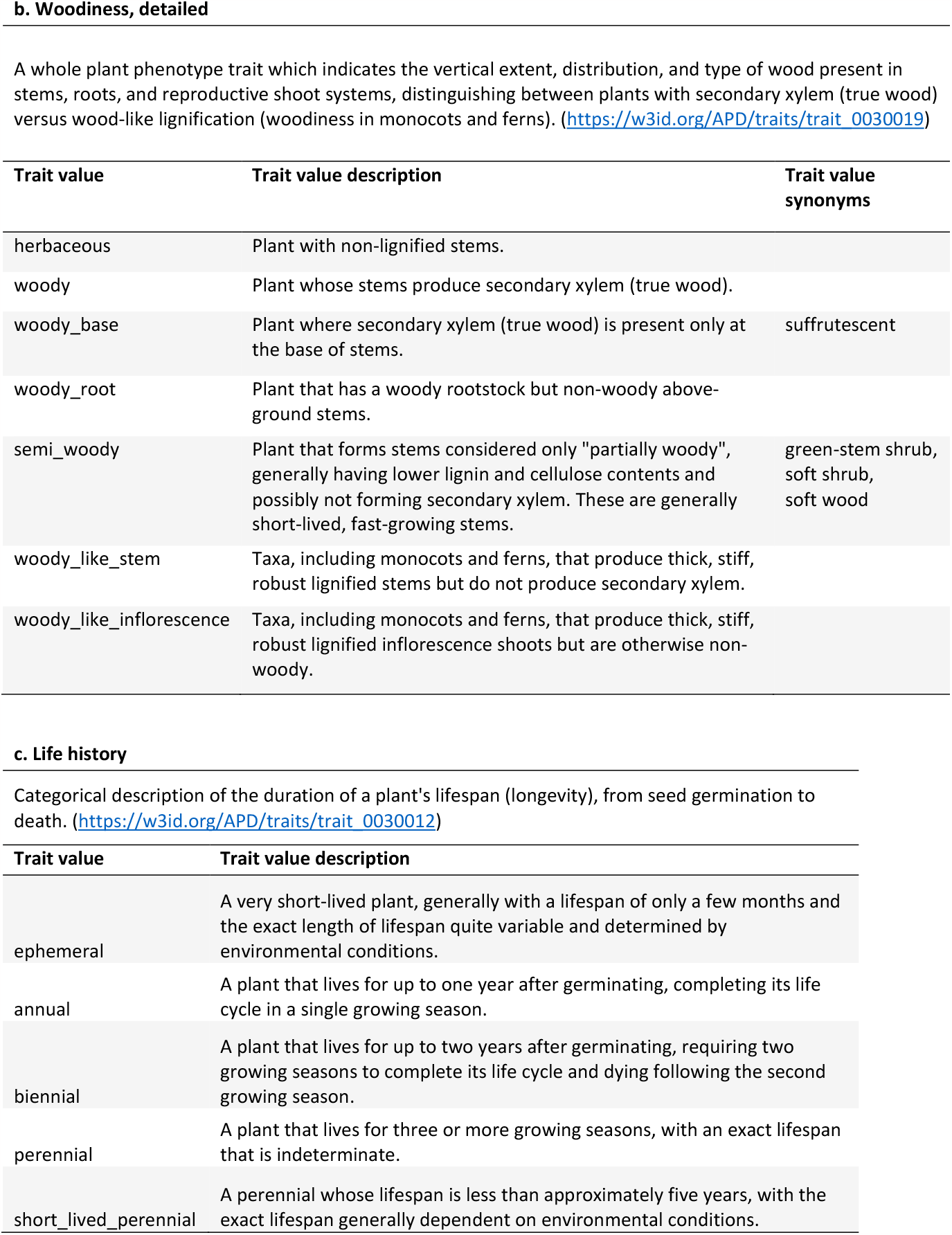
Definitions of traits and allowable values, used for scoring traits for each taxon. Definitions are sourced from the APD (Wenk *et al* 2023).

| Trait value | Trait value description | Trait value synonyms |
| --- | --- | --- |
| herb | A seed-bearing plant which does not have a woody above-ground stem; when available this term has been divided into more detailed terms. | forb |
| subshrub | A plant that is only woody at the base, with the remainder of the stems regenerating yearly. This term is not meant to encompass short-lived plants that are only borderline woody. | suffrutescent |
| shrub | A woody plant that is distinguished by NOT being a tree, due to one of more of the following characteristics, 1) they are less than 8 metres in height; 2) if a plant community includes trees, the shrubs are shorter than the surrounding trees and not part of the canopy; 3) in comparison to trees, the leaves are more distributed along the entire trunk; 4) the plant has multiple relatively narrow stems arising at or near ground level. The definition of 'shrub' is complex, as there are many single-stemmed shrubs within Australia and many taxa that are described in the taxonomic literature as a shrub or small tree. |  |
| mallee | Refers to Eucalyptus species which are small trees that form a large lignotuber. They have multiple stems arising at or near ground level and regenerate from the lignotuber following fire. |  |
| tree | A tall, woody, perennial plant, usually with a single main trunk, and its leaves predominantly elevated above the ground surface. |  |
| graminoid | Herbaceous plant with a grass-like morphology that is within the order Poales. Taxa both with and without a tussock form are mapped to this term. This term is most frequently applied to three large monocot families, Poaceae, Cyperaceae, and Juncaceae, but the term could apply to other taxa within Poales with a grass-like morphology, including narrow, linear leaves and non-showy flowers, including Restionaceae, Centrolepidaceae, Anarthriaceae, and Ecdeicoleaceae. Taxa outside Poales with a grass-like morphology should be designated as 'herbs' and also have the term 'tufted' mapped to the trait 'stem_growth_habit'. |  |
| tussock | A subcategory of graminoid, which is a plant where many shoots from the basal meristem form prominent tufts. |  |
| graminoid_not_tussock | A subcategory of graminoid, indicating a plant which explicitly lacks a tussock morphology, with the leaves not growing in tufts. |  |
| hummock | Plant that grows as a dense mound up to 1 metre in height. The term generally applies only to species within the genera *Triodia* that form such mounds and these are considered a distinctive and uniquely Australian growth form. |  |
| basal_large | A plant whose leaves form a distinctly large basal tuft or rosette, including large acaulescent monocots and cycads. |  |
| geophyte | Subcategory of herb, specifically referring to plants that survive the winter/dry season as an organ (rhizome, bulb, corm, or tuber) buried in dry soil, with all leaves dying back each year. |  |
| palmoid | Plant that bears a rosette-like canopy of typically large, often compound leaves atop a usually thick (pachycaulous), columnar, unbranched or little-branched stem. Palms, tree ferns, trunked cycads, trunked grass-trees, and *Pandanus* are mapped to this term. | rosette-tree, palm, grass-tree |
| climber | Plant that climbs up another plant's stem or branches, rather than being able to support itself. Term used only if datasets do not specify if a climbing plant is herbaceous (a vine) or woody (a liana). |  |
| climber_herbaceous | Herbaceous plant that climbs up another plant's stem or branches, rather than being able to support itself. This is usually synonymous with `vine`. Types of climbers (scrambling, twining) are captured under the trait `plant_climbing_mechanism`. | vine |
| climber_woody | Woody or thick-stemmed (>2 cm) plant that climbs up another plant's stem or branches, rather than being able to support itself. This is usually synonymous with `liana`. Woody climbers generally use hooks, tendrils, and/or adventitious roots to climb; the climbing mechanisms used by a taxon are captured under the trait `plant_climbing_mechanism`. | liana |
| fern | A taxonomic grouping; ferns are vascular plants that reproduce via spores. |  |
| lycophyte | A taxonomic grouping that includes quillworts, clubmosses, and selaginella. |  |
| herbaceous | Plant with non-lignified stems. |  |
| woody | Plant whose stems produce secondary xylem (true wood). |  |
| woody_base | Plant where secondary xylem (true wood) is present only at the base of stems. | suffrutescent |
| woody_root | Plant that has a woody rootstock but non-woody above-ground stems. |  |
| semi_woody | Plant that forms stems considered only "partially woody", generally having lower lignin and cellulose contents and possibly not forming secondary xylem. These are generally short-lived, fast-growing stems. | green-stem shrub, soft shrub, soft wood |
| woody_like_stem | Taxa, including monocots and ferns, that produce thick, stiff, robust lignified stems but do not produce secondary xylem. |  |
| woody_like_inflorescence | Taxa, including monocots and ferns, that produce thick, stiff, robust lignified inflorescence shoots but are otherwise non-woody. |  |

| Trait value | Trait value description |
| --- | --- |
| ephemeral | A very short-lived plant, generally with a lifespan of only a few months and the exact length of lifespan quite variable and determined by environmental conditions. |
| annual | A plant that lives for up to one year after germinating, completing its life cycle in a single growing season. |
| biennial | A plant that lives for up to two years after germinating, requiring two growing seasons to complete its life cycle and dying following the second growing season. |
| perennial | A plant that lives for three or more growing seasons, with an exact lifespan that is indeterminate. |
| short_lived_perennial | A perennial whose lifespan is less than approximately five years, with the exact lifespan generally dependent on environmental conditions. |

Automated trait extractions from Australia’s state and national floras by Coleman et al. (2023) formed the starting point for this work. They used Natural Language Processing (NLP; innovative text-string matching) to extract plant trait data for plant growth form, life history and woodiness from Australia’s national and state floras (Flora of Australia; PlantNet (NSW); VicFlora; FloraBase (WA); SA eFlora; and NT eFlora) and map them to the allowable trait values. In addition, trait values were inferred from higher-level taxa if, and only if, the higher-level taxon documented a single trait value. For instance, if the Flora of Australia indicated that a genus was comprised entirely of herbs, all taxa listed as species within the flora were scored as being herbs. For the state floras, only the taxa within that flora were scored by inference as the taxon descriptions may be written to just reflect the regional flora, not all Australian species; that is, a genus described within the Northern Territory flora of being comprised exclusively of herbs might include shrubs in Victoria (Coleman et al., 2003). Across all floras, Coleman et al. (2023) ultimately extracted plant growth form data for 25,396 taxa, life history data for 23,840 taxa and woodiness for 25,127 taxa (Table 2).

**Table 2.** Summary of how the traits were scored across taxa.

|  | trait |  |  |
| --- | --- | --- | --- |
|  | growth form | woodiness | life history |
| **automated scoring** |  |  |  |
| unambiguous scorings from floras | 20,784 | 24,084 | 22,314 |
| scored from growth form, woodiness | - | - | 3,045 |
| **manual scoring** |  |  |  |
| scored reading flora descriptions, including print volumes | 6,852 | 3,837 | 2,681 |
| primary taxonomic literature | 1,584 | 1,585 | 1,288 |
| reliable websites, non-Australian floras | 704 | 704 | 728 |
| AusTraits dataset* | 69 | 69 | - |
| **total taxa scored** |  |  |  |
| species and infraspecific taxa | 29,993 | 30,279 | 30,056 |
| species | 25,698 | 25,980 | 25,533 |

\* Mostly taxa known only by a phrase name that were scored in a dataset but for which no published taxon description exists.

As Tasmania does not have a comprehensive online flora that could be processed using NLP, we added trait data on Tasmanian plants manually transcribed by contributors to the Tasmanian print flora and submitted to AusTraits (Jordan 2001; McGlone et al. 2015). This combined dataset was filtered to exclude values that were either inconsistent across floras, had multiple values within a flora, or were designated as climbers. This left the following number of taxa with unique trait scores: plant growth form, 20,784; life history, 22,314; woodiness, 24,084. Hereafter, all listed methods refer to our refinement and expansion of this dataset.

To construct a dataset with a single harmonised trait value for all of Australia’s taxa, three additional steps beyond Coleman et al’s (2023) efforts were required: 1) manually interpreting ambiguous trait values; 2) merging disparate trait values offered by different floras; and 3) manually gap-filling trait data for all remaining taxa from the literature. The processes for completing each step are outlined below.

For each trait-by-taxon combination, if the starting compilation had unique, identical and unambiguous trait values across the floras, this value was accepted without review. This included all taxa with growth form consistently scored as either herbs, shrubs, trees, or graminoids across all floras that included the taxon. For life history, this included all taxa that were scored consistently as annual, biennial, or perennial. For woodiness, all taxa scored consistently as either ‘woody’ or ‘herbaceous’ were accepted. Taxa that were reported as possessing multiple trait values, either within the same source or between multiple sources, were not automatically accepted. This included taxa that were scored as being either annual or perennial within a single flora, scored in AusTraits as ‘annual perennial’, and taxa with inconsistent scorings across floras. For these taxa, the original descriptions in each flora were read to ensure it was appropriate to include multiple trait values for that species. The individual taxon descriptions were also read for all climbing plants, as the floras use a diverse vocabulary to describe climbers and automated extractions were inconsistently able to identify a climber as woody versus herbaceous.

For taxa without information in the floras, additional resources were relied upon, including the taxonomic literature, print floras, online floras for regions outside Australia, and websites deemed to be reliable. As the state of Queensland lacks a comprehensive flora and the Northern Territory online flora is incomplete, most plants requiring manual scoring had Australian ranges restricted to these states. Additionally, taxa described in the last decade have generally not yet been added to the online state and national floras. Descriptions for naturalised species were, when available, sourced from an online flora for their native range. Additionally, some online compilations were deemed to be reliable sources of information. For instance, Wikipedia has an increasing number of taxon descriptions as its taxon descriptions are linked to iNaturalist (“iNaturalist” 2023). Useful Tropical Plants (Fern and Fern 2023b) and Useful Temperate Plants (Fern and Fern 2023a) offer well-curated descriptions of both tropical and temperate plants with known human uses. In addition, for invasive naturalised species, there are many Australian and global weed databases. For all these sources, trait values were manually scored. Plant growth form and woodiness were scored first, with life history trait values scored during a second data collection campaign. The same references and process were used each time, but slightly different numbers of trait values were extracted between the two initiatives. For many taxa, life history could be scored based on the plant growth form and woodiness scores, as any shrub or tree was mapped as perennial and most woody plants could be mapped as perennial.

Among the taxa with trait values that were not automatically accepted from Coleman et al. (2023) or which lacked scores in Coleman et al., most were scored by manually reading the taxon descriptions in the either the various Australian online floras, Flora of Australia print volumes, or the Flora of Southeast Queensland (Table 2). In many cases this simply required looking at the extracted phrase and confirming multiple trait values within a single flora or disparate scorings across floras were accurate and appropriate. The taxonomic literature added a further 1584 plant growth form values, 1585 woodiness values, and 1288 life history values (Table 2). Websites, including online floras for regions outside Australia, were the source of information for 704 plant growth form and woodiness scores and 728 life history scores (Table 2).

In total, we report plant growth form values for 29,993 taxa, woodiness values for 30,279 taxa, and life history scores for 30,056 taxa; all but a few hundred of the 30,494 taxa in the initial list of APC-accepted taxa were scored (Table 3). While most taxa display a unique value for each of the three traits, a significant component of Australia’s flora is polymorphic with respect to plant growth form, life history, or life history, with multiple trait values appropriate for 4871 taxa for one or more traits.

**Table 3.** Frequency of trait values across taxa for three traits. Multiple values are taxa with two or more trait values recorded. A single taxon will therefore appear in more than one row in that column.

| trait value | Number of taxa |  |
| --- | --- | --- |
|  | unique value | multiple value |
| **Plant growth form** |  |  |
| herb | 9095 | 792 |
| shrub | 10170 | 2640 |
| tree | 2265 | 2193 |
| graminoid | 1938 | 2 |
| tussock | 918 | 0 |
| fern | 476 | 28 |
| climber_woody | 410 | 171 |
| subshrub | 327 | 602 |
| climber_herbaceous | 302 | 169 |
| mallee | 281 | 294 |
| palmoid | 139 | 38 |
| basal_large | 73 | 12 |
| hummock | 65 | 0 |
| lycophyte | 51 | 0 |
| geophyte | 25 | 0 |
| climber | 0 | 7 |
| **Total** | **26535** | **3446** |
| **Woodiness** |  |  |
| herbaceous | 12767 | 911 |
| woody | 16149 | 389 |
| semi_woody | 208 | 240 |
| woody_base | 66 | 407 |
| woody_like_stem | 78 | 0 |
| woody_root | 2 | 49 |
| **Total** | **29270** | **1074** |
| **Life history** |  |  |
| perennial | 25616 | 820 |
| annual | 2857 | 1165 |
| biennial | 40 | 246 |
| short_lived_perennial | 29 | 373 |
| ephemeral | 11 | 93 |
| **Total** | **28553** | **1250** |

Data for plant growth form and woodiness are included in AusTraits under the dataset_id Wenk_2022 (from v4.1.0) and those for life history under the dataset_id Wenk_2023 (from v4.2.0) and are most easily extracted using the {austraits} R package (Falster et al. 2021).

The following R code can be used to access the complete dataset for plant growth form, woodiness, and life history:

~~~
remotes::install_github(“traitecoevo/austraits”, dependencies = TRUE, upgrade = “ask”) austraits <-austraits::load_austraits(doi = “10.5281/zenodo.10156222”)
complete_traits <-austraits$traits %>%
  dplyr::filter(dataset_id %in% c(“Wenk_2022”, “Wenk_2023”)) %>%
  dplyr::select(taxon_name, trait_name, value) %>%
  dplyr::distinct(taxon_name, trait_name, .keep_all = TRUE) %>%
  tidyr::pivot_wider(names_from = trait_name, values_from = value)
~~~

For further information of accessing and manipulating AusTraits data, see: https://traitecoevo.github.io/traits.build-book/AusTraits_tutorial.html

## Discussion

In recognition of the need for more complete tabular trait data for Australian plants, we present the first near-complete continental-scale scoring of a trio of categorical traits. This dataset is a resource ready for use for diverse research and management programs. The data can be used to explore trait distributions across climatic gradients, as a covariate in ecological analyses, and offers a mechanism to rapidly assess species’ responses to natural disasters or to inform long-term management decisions. Analyses have indicated that plants with different values for the traits in question (life history, growth form, and stem woodiness) exhibit different functional trade-offs (Šímová et al. 2018; Vico et al. 2016; Towers et al. 2023). Similarly, plants with different values for these three categorical traits display different numeric trait-climate associations (Flores-Moreno et al. 2019), require different conservation strategies (Mayfield, Ackerly, and Daily 2006), have different responses to key disturbances (Gallagher et al. 2021; Clarke et al. 2013), and even differ in the degree to which they have a public photographic record (Mesaglio et al. 2023). Having complete tabular datasets for these traits across the Australian flora, therefore, has direct implications for the management and conservation of these species, in addition to facilitating more broad-scale biogeographic analyses.

Increasing awareness of the existence of this resource should accelerate its use in research and eliminate the need for other researchers to unnecessarily expend time and resources creating their own datasets for these traits. For example, 102 of the 363 datasets within the most recent AusTraits release included the trait plant growth form. This occasionally represents values scored in the field, but mostly these are data transcribed from floras and added into an analysis as a descriptive or statistical covariate. This information and data on life history and woodiness can now be quickly sourced from the compilation presented here. Many additional studies will have chosen study taxa based on their scores for plant growth form, life history, and woodiness; this dataset represents a rapid way to scan a list of potential study taxa for appropriate study taxa. Within our research group, the plant growth form dataset has already been used twice (Towers et al. 2023; Mesaglio et al. 2023).

The need to manually score or check the scores of these three traits for more than 25% of Australia’s plant taxa highlights the need for comprehensive, open-access, tabular resources for Australia’s flora. Most taxa missing trait values were those restricted (within their Australian range) to Queensland, the Northern Territory or offshore islands, as these regions lack comprehensive online floras. For instance, of the 162 described taxa lacking life history scores, only 13 have ranges that include New South Wales, only 6 have ranges that include Victoria, while 82 have ranges that include Queensland and 55 have ranges that include the Northern Territory. The eventual completion of the Flora of Australia will alleviate this problem, but as long as this resource is incomplete, trait data about taxa restricted to Australia’s northern regions remains difficult to compile.

We suggest that new taxon descriptions and taxon revisions should explicitly include data for these traits in a standardised format, even if it is only to acknowledge that a taxon displays a range of trait values. A taxon may be rare and known from only a few localities, but its conservation is impeded if its annual or perennial status is unknown (Gallagher et al. 2021; Butt and Gallagher 2018). Another concern is that new taxon descriptions may be presented in books or hard-to-access taxonomic journals, instead of the generally open-access journals associated with national or state herbaria. For instance, many species of *Drosera* and *Eremophila* are only described in books (Lowrie 2013; Chinnock 2007). Until the Flora of Australia adds profiles for all taxa, portals like Wikipedia and World Flora Online are the main online resources (e.g. the profiles for *Eremophila* species have all been transcribed from Chinnock 2007 into Wikipedia). Although Plants of the World Online includes plant growth form and life history data for most taxa, its descriptions are not always correct and were sometimes rejected as references. For instance, several instances of the nonsensical phrase “climbing herbaceous trees” were noted (e.g. for *Merremia gemelli*), suggesting an automatic extraction of words suggesting a vine that climbed into the treetops.

While it is fantastic to be able to offer a resource with trait values for 99% of Australia’s flora, it is simultaneously disappointing that these core traits were not scorable for all described taxa. For instance, there does not exist a reference that unequivocally states whether *Zornia oligantha* is annual, perennial, or displays a polymorphic life history. Taxa that could not be scored either had not had a taxon revision since 1950 (and usually since 1920) or the taxon description from the taxonomic literature did not include information on the three traits being scored. References from before 1950 could only haphazardly be relied upon as most of these older descriptions, if accessible, rarely included information on life history for herbaceous species and did not consistently distinguish between herbs and subshrubs (and hence, taxon woodiness). There also exist a small number of genera where the most recent taxon revisions incompletely documented these traits, generally because species’ phenotypes span the discrete trait values or display an intermediate, difficult-to-express trait value. Three genera within which individual taxa were rarely scored for life history, *Eriocaulon, Nicotiana* and *Spermacoce*, can likely grow and reproduce as annuals or perennials, depending on the environmental conditions. Many species in the genera *Sida* and *Solanum* are difficult to score for plant growth form and woodiness, as many taxa are short-lived, post-disturbance species that grow rapidly, assuming a tall, well-branched form, but have only partially lignified stems. The phrases ‘annual shrub’ (*Roepera reticulata* in WA’s Florabase) and ‘soft shrub’ (*Grevillea fililoba* in Flora of Australia) exemplify these ‘intermediate’ forms. The woodiness trait value ‘semi_woody’ is intended for such instances but can only be used if the taxon description includes terminology to suggest a partially lignified stem.

We hope the existence of these datasets encourages researchers to use the data to explore evolutionary and ecological questions about Australia’s diverse flora as well as to use these data as co-variates in other trait-based analyses. We additionally hope that the compilation and release of this dataset will not just offer new research opportunities, but also inspire other researchers to work toward gap-filling other essential plant traits and sharing their results on open-source, harmonised data platforms, like AusTraits.

## Acknowledgements

We acknowledge the work of all Australian taxonomists and their supporting institutions, whose long-term work on writing flora descriptions has enabled this work: Australian National Botanic Gardens; Australian National Herbarium; Biodiversity Science, Parks Australia; Centre for Australian National Biodiversity Research; Department of Biodiversity, Conservation and Attractions, Western Australia; Department of Environment, Land, Water and Planning, Victoria; Flora of Australia; National Herbarium of NSW; National Herbarium of Victoria; Northern Territory Herbarium; Queensland Herbarium; State Herbarium of South Australia; Tasmanian Herbarium; and the Western Australian Herbarium. We thank Russell Barrett, Ashleigh Ford, Hervé Sauquet, Guy Taseski, Sophie Yang for assistance with the manuscript, scoring of taxa, and ideas about how best to fill gaps in the dataset.

